# Hexosamine pathway activation improves memory but does not extend lifespan in mice

**DOI:** 10.1101/2022.01.07.475325

**Authors:** Kira Allmeroth, Matías D. Hartman, Martin Purrio, Andrea Mesaros, Martin S. Denzel

## Abstract

Glucosamine feeding and genetic activation of the hexosamine biosynthetic pathway (HBP) have been linked to improved protein quality control and lifespan extension in various species. Thus, there is considerable interest in the potential health benefits of dietary supplementation with glucosamine or other HBP metabolites in people. The HBP is a sensor for energy availability and its activation has been implicated in tumor progression and diabetes in higher organisms. As the activation of the HBP has been linked to longevity in lower animals, it is imperative to explore the long-term effects of chronic HBP activation in mammals, which has not been examined so far. To address this issue, we activated the HBP in mice both genetically and through metabolite supplementation, and evaluated metabolism, memory, and survival. GlcNAc supplementation in the drinking water had no adverse effect on weight gain in males but increased weight in young female mice. Glucose or insulin tolerance were not affected up to 20 months of age. Of note, we observed improved memory in the Morris water maze in young male mice supplemented with GlcNAc. Survival was not changed by GlcNAc supplementation. To assess the effects of genetic HBP activation we overexpressed the key enzyme GFAT1 as well as a constitutively activated point mutant form in all mouse tissues. We detected elevated UDP-GlcNAc levels in mouse brains, but did not find any effects on behavior, memory, or survival. Together, while dietary GlcNAc supplementation did not extend survival in mice, it positively affected memory and is generally well tolerated.

## Introduction

The hexosamine biosynthetic pathway (HBP) is an anabolic pathway that consumes fructose-6-phosphate (Fru6P), glutamine, acetyl-CoA, and UTP to produce the high energy molecule uridine 5’-diphospho-N-acetyl-D-glucosamine (UDP-GlcNAc) (Figure 1a). The HBP thus requires carbohydrate, aminoacidic, lipidic, and nucleotide donors and is centrally positioned at a cross-roads of energy metabolism (Wells, Vosseller, and Hart 2003). The first step in the HBP is catalyzed by glutamine fructose- 6-phosphate amidotransferase (GFAT), that employs L-glutamine (Gln) as a nitrogen donor to convert Fru6P to D-glucosamine-6-phosphate (GlcN6P), diverting 2-3% of cellular glucose into the HBP (Marshall, Bacote, and Traxinger 1991). Next, glucosamine 6-phosphate N-acetyltransferase (GNA1) catalyzes the acetylation of GlcN6P using acetyl-CoA to produce N-acetyl-D-glucosamine-6-phosphate (GlcNAc6P); the phosphoglucomutase PGM3 then isomerizes GlcNAc6P to yield GlcNAc1P and finally, the phosphorylase UAP1 uses UTP to form UDP-GlcNAc. UDP-GlcNAc and its epimer UDP-GalNAc are precursors for biopolymer synthesis and for distinct glycosylation reactions including N- and O-linked glycosylation, and O-GlcNAcylation (Hanisch 2001; Hart 1997; Parodi 2000).

**Figure 1:**
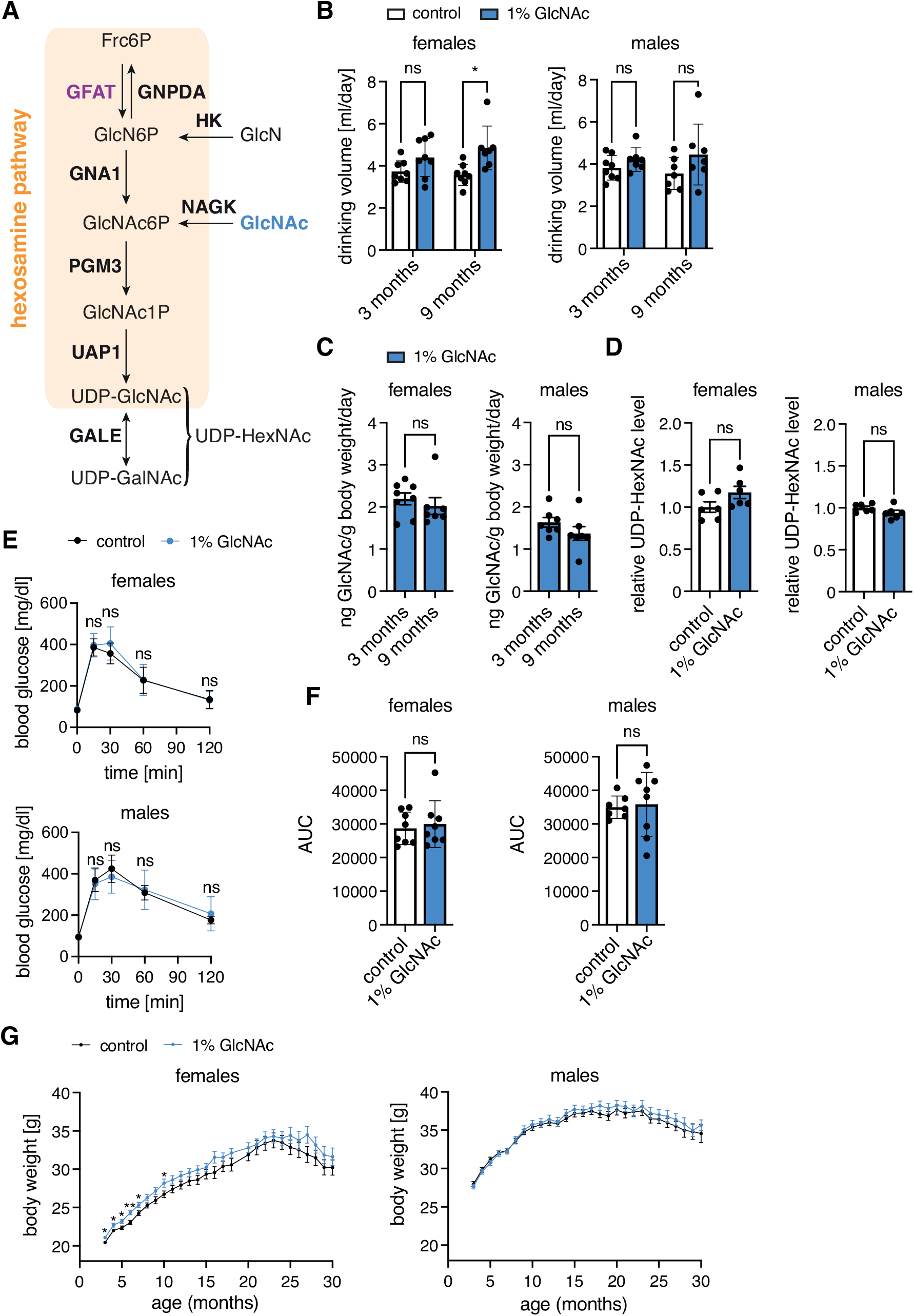
Hexosamine biosynthetic pathway activation by GlcNAc feeding does not have adverse effects in mice. (A) Schematic representation of the hexosamine biosynthetic pathway. The rate-limiting enzyme GFAT is depicted in purple, GlcNAc is marked in blue. (B) Drinking volume of control (white) and GlcNAc-treated mice (blue) of both sexes at 3 and 9 months of age. Data are presented as mean ± SD (n≥7). Two-way ANOVA, Tukey’s post-test; * p<0.05; ns: not significant. (C) GlcNAc consumption in ng/g body weight per day of mice of both sexes at 3 and 9 months of age. Data are presented as mean ± SD (n≥7). Unpaired t-test; ns: not significant. (D) Relative UDP-HexNAc levels in hemibrains of control (white) and GlcNAc-treated mice (blue) of both sexes. Data are presented as mean ± SEM (n=6). Unpaired t-test; ns: not significant. (E) Blood glucose concentration at 0 (fasting), 15, 30, 60, and 120 min after intraperitoneal injection of glucose solution (2 g/kg body weight) of control (black) and GlcNAc-treated mice (blue) of both sexes at 10 months of age. Data are presented as mean ± SD (n≥7). Multiple unpaired t-tests; ns: not significant (F) Area under the curve (AUC) calculated using data shown in (E). Data are presented as mean ± SD (n≥7). Unpaired t-test; ns: not significant (G) Body weight of control (black) and GlcNAc-treated mice (blue) of both sexes from 3 to 30 months of age. Data are presented as mean ± SD (n≥13). Multiple unpaired t-tests; ** p<0.01; * p<0.05; only significant changes are indicated.

GFAT is the HBP’s key enzyme and it is composed of two domains: the glutaminase domain that hydrolyses L-Gln to produce ammonia and L-glutamate (L-Glu), and the isomerase/transferase domain, which isomerizes Fru6P into Glc6P and catalyzes the transfer of ammonia to produce GlcN6P (Denisot, Goffic, and Badet 1991). In eukaryotes, GFAT1 is feedback inhibited by the end product of the HBP, UDP-GlcNAc. Our previous observations have delineated a number of single amino acid substitutions in GFAT1 that lead to increased activity through loss of UDP-GlcNAc feedback inhibition and altered domain interactions, resulting in elevated HBP flux and cellular UDP-GlcNAc accumulation (Denzel et al. 2014; Horn et al. 2020; Ruegenberg et al. 2020, 2021). Two alternative reactions feed into the HBP and increase the flux, bypassing the finely regulated GFAT-mediated step: glucosamine (GlcN) can be incorporated through its phosphorylation via hexokinase to generate GlcN6P (Stocchi et al. 1981), whereas N-acetyl-D-glucosamine (GlcNAc) can be incorporated through its phosphorylation via N-acetyl-D-glucosamine kinase producing GlcNAc6P (Weihofen et al. 2006). Overall, HBP activation can be achieved through the overexpression of the rate-limiting step enzyme GFAT1 or through the supplementation of the metabolites GlcN or GlcNAc.

O-GlcNAcylation is one mechanism linking UDP-GlcNAc availability, hence HBP activity, to protein function through post-translational modifications. Hyper-O-GlcNAcylation is observed in many cancer types, suggesting that this modification is a key molecular event in tumor formation, progression, aggressiveness, and potentially a new cancer hallmark (Fardini et al. 2013). Elevated GFAT1 expression has been associated with poor overall survival in patients suffering hepatocellular carcinoma (Li et al. 2017) and, strikingly, GFAT inhibitors like the diazo-derivative of serine, azaserine, and 6-diazo-5-oxo-L-norleucine (DON) decrease HBP flux and exhibit anti-tumor activity (Lemberg et al. 2018). The HBP has also been shown to be involved in diabetes; HBP activation in young rats by GlcN infusion leads to insulin resistance and enhanced transcription factor glycosylation, resembling an aging phenotype (Einstein et al. 2008; Marshall, Bacote, and Traxinger 1991). Moreover, GlcN impairs the GLUT4 glucose transporter (whose expression is stimulated by insulin), resembling an insulin-resistant state (Baron et al. 1995). These data highlight the importance of monitoring potential adverse side effects upon chronic HBP activation.

Although high HBP flux has been linked to cancer and diabetes, previous observations also support a plausible role in longevity and prevention of age-related pathologies (Denzel and Antebi 2015). The HBP is involved in protein quality control and its activation leads to improved protein homeostasis and lifespan extension in *Caenorhabditis elegans*: worms with GFAT-1 gain-of-function (GOF) mutations have increased UDP-GlcNAc/UDP-GalNAc levels and are long-lived (Denzel et al. 2014). Strikingly, this lifespan extension is recapitulated by GlcNAc feeding in a dose-dependent manner (Denzel et al. 2014). Elevating HBP metabolite levels through GFAT-1 GOF mutations or through GlcNAc supplementation enhances ER protein quality control, ER-associated protein degradation (ERAD), and autophagy (Denzel et al. 2014; Shintani, Kosuge, and Ashida 2018). Of note, worms overexpressing WT GFAT-1 or supplemented with GlcNAc activate the integrated stress response (ISR) (Horn et al. 2020). In mammalian Neuro-2a (N2a) cells, mild HBP activation through GFAT1 GOF mutation or through GlcNAc feeding likewise induces the ISR: the PERK branch is activated leading to increased phosphorylation of the α subunit of eukaryotic initiation factor 2 (eIF2α) and upregulation of the stress master transcription factor ATF4 (Horn et al. 2020). In N2a cells, HBP activation has a protective role against proteotoxicity in a PERK- and autophagy-dependent manner and muscle cell-specific HBP activation rescues polyQ-mediated toxicity in worms, a process that depends on ATF4 (Horn et al. 2020). GlcN supplementation likewise induces ER stress in rat cells (Lombardi et al. 2012), and it leads to eIF2α phosphorylation through the PERK branch with a concomitant mRNA translation arrest (Kline et al. 2006). Overall, these data suggest a protective role of HBP activation in various species.

GlcN feeding extends lifespan in worms and mice, in a manner that is independent from the HBP: GlcN supplementation impairs glucose metabolism and activates AMPK to promote mitochondrial biogenesis, which mimics low-carbohydrate diet paradigms (Weimer et al. 2014). GlcN supplementation in mice leads to glycolysis impairment as a rise in GlcN6P was observed (Weimer et al. 2014), a metabolite that inhibits hexokinase (Silverman 1963). Strikingly, GlcN supplementation does not increase the levels of UDP-GlcNAc in the mouse liver. Furthermore, silencing PGM3 within the HBP by RNAi does not impair GlcN-mediated lifespan extension in worms (Weimer et al. 2014). Instead, GlcN treatment induces a metabolic switch towards increased amino acid catabolism in mice (Weimer et al. 2014), supporting the HBP- independent effect. Given the various observed effects of GlcN metabolism and HBP activation, their role in mammalian life- and healthspan needs to be explored.

To investigate a possible modulatory role of the HBP in mammalian aging, we employed distinct strategies to increase HBP activity and examined general behavior, memory, and survival in mice. We found that GlcNAc supplementation in the drinking water or overexpression of wildtype or GOF GFAT1 did not have adverse effects in mice. Importantly, the interventions did not interfere with glucose metabolism. Mice supplemented with dietary GlcNAc or overexpressing GFAT1 did not exhibit lifespan extension. Nevertheless, we observed memory improvement in young male mice fed with GlcNAc, suggesting a beneficial effect of HBP activation in mammals.

## Results

### HBP pathway activation by GlcNAc feeding does not have adverse effects in mice

To chronically activate the HBP, we supplemented the drinking water of C57BL/6J mice with 1% GlcNAc starting at 8 weeks of age. We first tested whether GlcNAc feeding might lead to adverse reactions. Regarding water intake, females consumed more GlcNAc-supplemented water compared to controls at 9 months of age; males, however, did not show any difference in water consumption (Figure 1b). The drinking volume of around 4 ml per day corresponds to a GlcNAc consumption of 1.5-2 ng*g^-1^ body weight per day (Figure 1c). Importantly, GlcNAc supplementation in the drinking water had no effect on food intake (Figure S1a). We used LC-MS to quantify UDP-HexNAc levels (combined UDP-GlcNAc and UDP-GalNAc) in brain lysates from 9 months old mice to test effects of dietary supplementation. Surprisingly, after 7 months of GlcNAc feeding, the UDP-HexNAc levels were not affected in mouse brains (Figure 1d); similarly, liver samples did not show increased levels of UDP-HexNAc (data not shown). Because elevated flux through the HBP has been associated with insulin resistance and diabetic complications (Han, Chen, and Holloszy 2003; Rossetti et al. 1995), we tested the effect of chronic GlcNAc supplementation on glucose utilization by measuring blood glucose clearance and insulin-stimulated glucose utilization. Chronic GlcNAc intake had no detectable effect on either blood glucose clearance or insulin response in 10 months old mice (Figure 1e,f and Figure S1c), or 20 months old mice (Figure S1b,c). In 4 months old females, however, we observed slightly higher blood glucose levels 30 and 60 min after the injection of insulin (Figure S1c). Nevertheless, overall, we conclude that GlcNAc supplementation did not alter insulin signaling. While males did not display any difference in body weight, GlcNAc-supplemented female mice showed increased body weight compared to controls (Figure 1g). Indirect calorimetric measurements revealed an increased respiratory exchange ratio (RER) in 9 months old female mice during the day, while no difference was detected in age-matched male mice (Figure S1d). This increased RER, as well as the effects on body weight, might be caused by the elevated GlcNAc consumption (Figure 1b,c), which was also sex-specific. Despite these effects, female mice did not display changes in insulin signaling after 4 months. Therefore, overall, our data suggest that chronic GlcNAc supplementation does not have negative side effects in mice.

### GlcNAc supplementation does not affect coordination or neuromuscular function

Having excluded negative side effects of dietary GlcNAc supplementation on the metabolic health of mice, we next aimed to test possible effects of GlcNAc supplementation on coordination and basic neuromuscular function. To this end, the fitness of mice at 6 months of age was analyzed using the rotarod, and by assessing grip strength and treadmill endurance. Of note, GlcNAc supplementation did not affect grip strength (Figure 2a-b), rotarod performance (Figure 2c), or forced maximal endurance on a treadmill (Figure 2d). Thus, fitness and locomotion of mice were not changed by GlcNAc supplementation.

**Figure 2:**
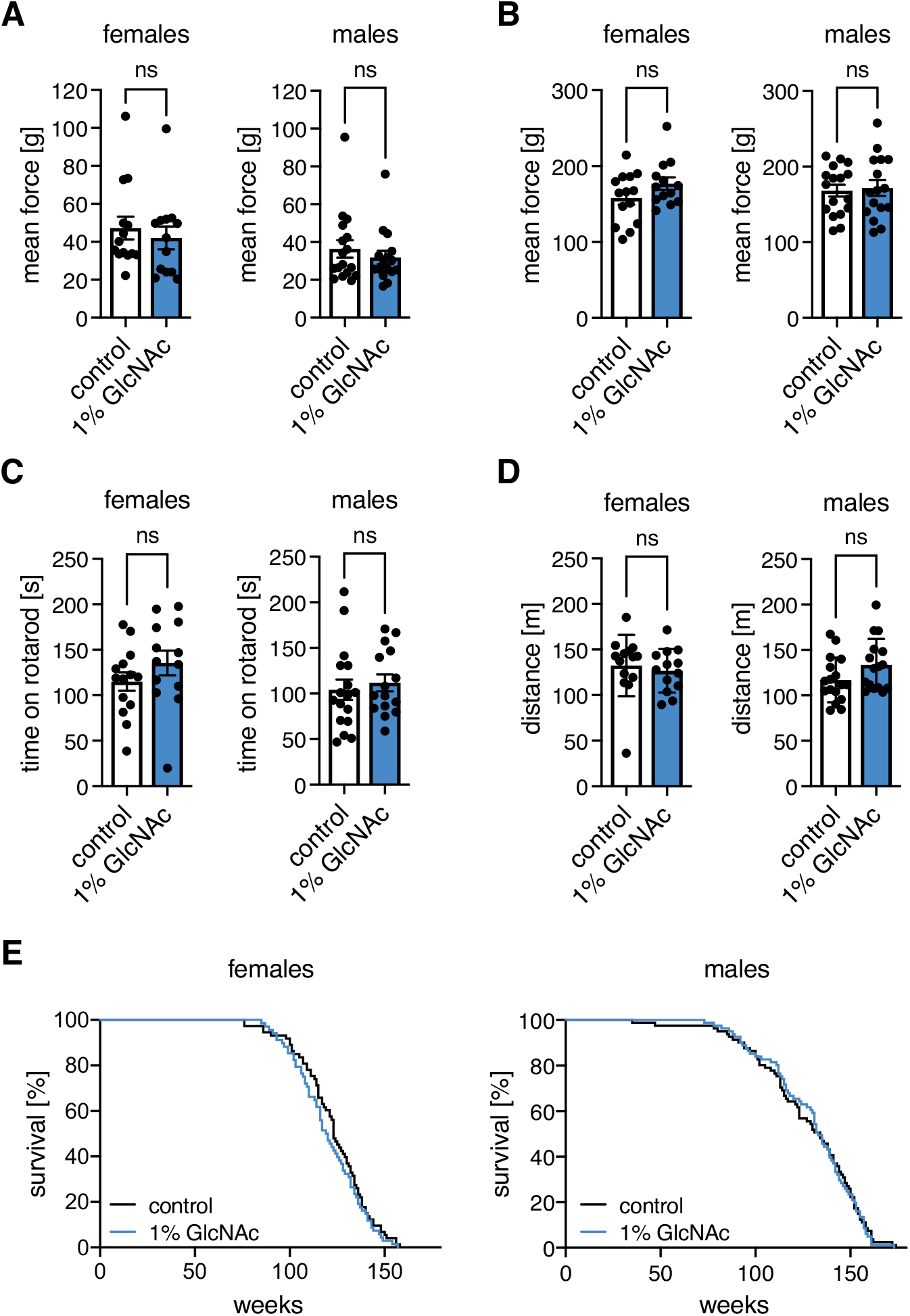
GlcNAc supplementation does not influence fitness of mice. (A) Mean force measured in a grip strength test with two paws of control (white) and GlcNAc- treated mice (blue) of both sexes at 6 months of age. (B) Mean force measured in a grip strength test with four paws of control (white) and GlcNAc-treated mice (blue) of both sexes at 6 months of age. (C) Maximal time on a rotarod of control (white) and GlcNAc-treated mice (blue) of both sexes at 6 months of age. (A-C) Data are presented as mean ± SEM (n≥13). (D) Maximal distance on a treadmill of control (white) and GlcNAc-treated mice (blue) of both sexes from at 6 months of age. Data are presented as mean ± SD (n≥13). (A-D) Unpaired t-test; ns: not significant (E) Lifespan analysis of control (black) and GlcNAc-treated mice (blue) of both sexes (females: n=69; males: n=82).

As GlcNAc supplementation extends *C. elegans* lifespan (Denzel et al. 2014) and showed no adverse effects in mice, we supplemented mice of both sexes with GlcNAc in the drinking water from week 8 until they died to assess the effect on mammalian lifespan. GlcNAc feeding did not extend mouse survival (Figure 2e). Importantly however, survival was not reduced by GlcNAc supplementation. In sum, these data support that chronic HBP activation by GlcNAc feeding does not have adverse side effects on general fitness and health in mice.

### GlcNAc feeding improves memory of young male mice

We next aimed to test memory and spatial cognition in the Morris water maze (Vorhees and Williams 2006) in mice aged 4 months. To exclude that the results were influenced by changes in locomotion or behavior, we first assessed general activity as well as exploratory behavior in the open field test. GlcNAc feeding did not alter spontaneous locomotor activity or exploratory behavior as measured by the distance, speed, and percentage of distance spent in the center of the open field at 6 months of age (Figure 3a-b; Figure S2a). Additionally, analysis of the home cage activity in metabolic cages did not reveal differences caused by GlcNAc supplementation (Figure S2b). Thus, locomotion and exploratory behavior were not affected by GlcNAc treatment.

**Figure 3:**
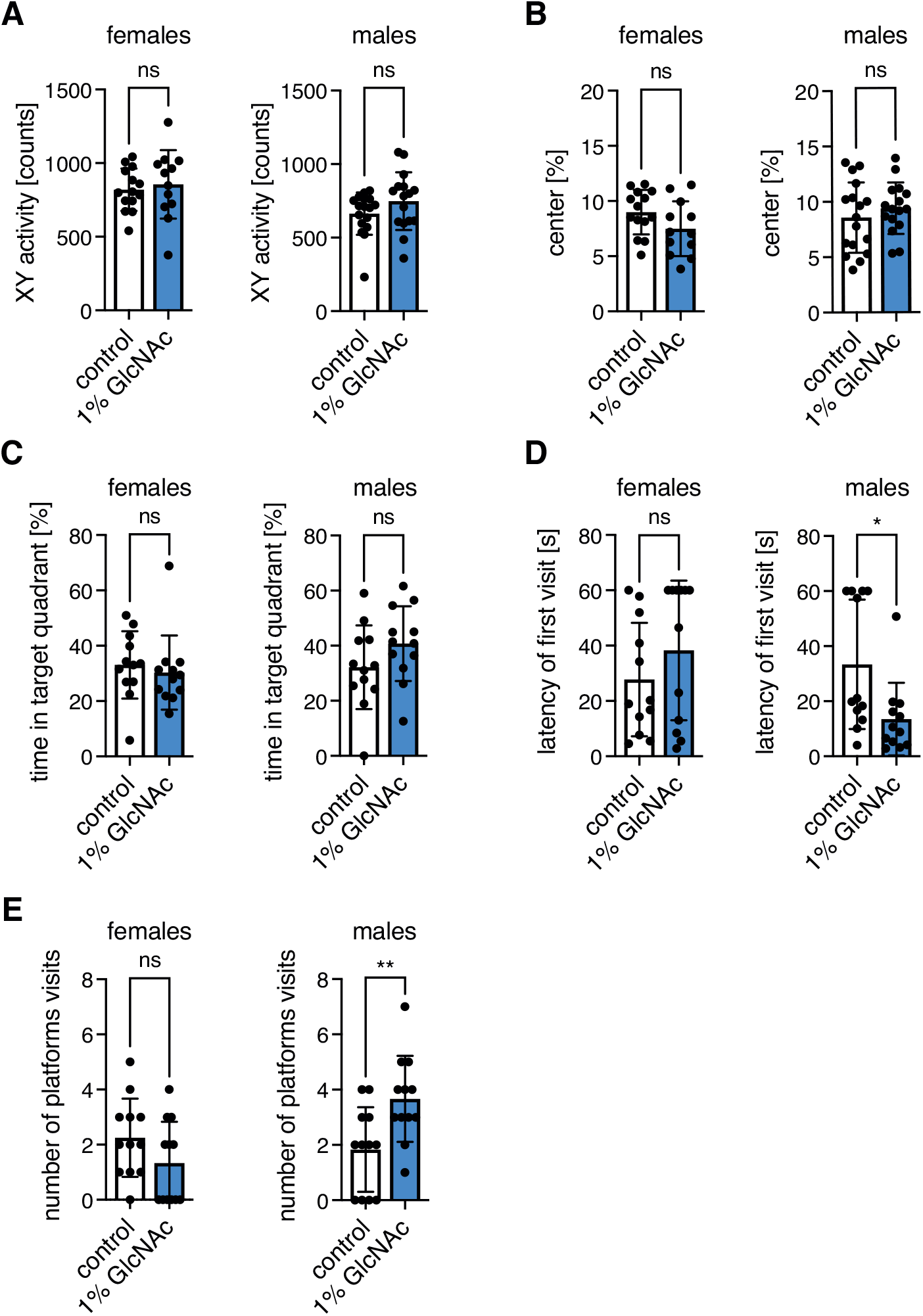
GlcNAc feeding improves memory of young male mice. (A) XY activity measured in the open field test of control (white) and GlcNAc-treated mice (blue) of both sexes at 6 months of age. (B) Percent of distance spent in the center of the open field of control (white) and GlcNAc-treated mice (blue) of both sexes at 6 months of age. (A-B) Data are presented as mean ± SD (n≥12). (C) Percent of time spend in the target quadrant, (D) Latency of the first platform visit, and (E) Number of platform visits upon removal of the hidden platform in the Morris water maze test of control (white) and GlcNAc-treated mice (blue) of both sexes at 4 months of age. (C-E) Data are presented as mean ± SD (n=12). (A-E) Unpaired t-test; ** p<0.01; * p<0.05; ns: not significant.

In the Morris water maze, the animals were trained for consecutive 5 days, and after the last session, memory was assessed by removal of the hidden platform. During training, both distance and searching time (latency) decreased progressively (Figure S2c-d), while swimming speed remained constant (Figure S2e). These data suggest that the mice learned to find the hidden platform. There was no significant difference between GlcNAc-fed animals compared to controls (Figure S2c-e) during the training period. Still, GlcNAc-supplemented male mice tended to swim shorter distances and to spend less time in the water from day 2 on (Figure S2c-d). During the test, there was no difference in the time mice spent in the target quadrant upon removal of the platform (Figure 3c), strikingly, however, the latency of the first platform visit (defined as the time needed until the mice cross the area where the platform used to be) was strongly decreased in GlcNAc-fed males (Figure 3d). Furthermore, the number of platform visits (defined as the number of times the mice cross the area where the platform used to be) was increased in GlcNAc-fed male mice (Figure 3e). Overall, our observations suggest that GlcNAc feeding improves memory of young male mice.

### HBP activation by overexpression of WT or G451E huGFAT1 does not influence body weight in mice

To corroborate the data obtained upon dietary GlcNAc supplementation, we pursued a parallel approach to genetically activate the HBP by overexpressing N-terminally FLAG-HA tagged human GFAT1 (huGFAT1) in all mouse tissues. The *Rosa26* locus was engineered to contain an expression construct composed of a loxP-flanked transcription termination cassette upstream of the huGFAT1 open reading frame (huGFAT1 wt tg^+/-^, Figure 4a). These mice were crossed with transgenic CMV-cre^+/-^ females, to obtain huGFAT1 overexpressing animals (huGFAT1 wt OE, Figure 4a). The functionality of the cassette was confirmed by Western blot analysis (Figure 4b): In fibroblasts isolated from huGFAT1 wt tg^+/-^ newborn mice HA expression was not detectable due to the lack of cre recombinase expression. Endogenous GFAT1 expression was comparable to WT and CMW-cre^+/-^ fibroblasts. huGFAT1 wt OE fibroblasts in which the GFAT1 transgene and cre recombinase were co-expressed, displayed elevated GFAT1 and HA expression, demonstrating successful expression of the transgene. Accordingly, UDP-HexNAc levels were comparable in WT, huGFAT1 wt tg^+/-^ and CMV-cre^+/-^ fibroblasts and about 2-fold elevated by GlcNAc supplementation and huGFAT1 wt OE (Figure 4c). As expected, huGFAT1 overexpression also resulted in elevated levels of UDP-GlcNAc (Figure 4d) and UDP-GalNAc (Figure S3a) in the brains of 3 months old male and female mice compared to brains of WT and CMV-cre^+/-^ mice. Body weight analysis over 27 months indicated that there was no effect of huGFAT1 wt overexpression compared to CMV-cre^+/-^ mice in both sexes (Figure 4e). However, cre expression slightly reduced body weight in male mice compared to WT controls (Figure 4e).

**Figure 4:**
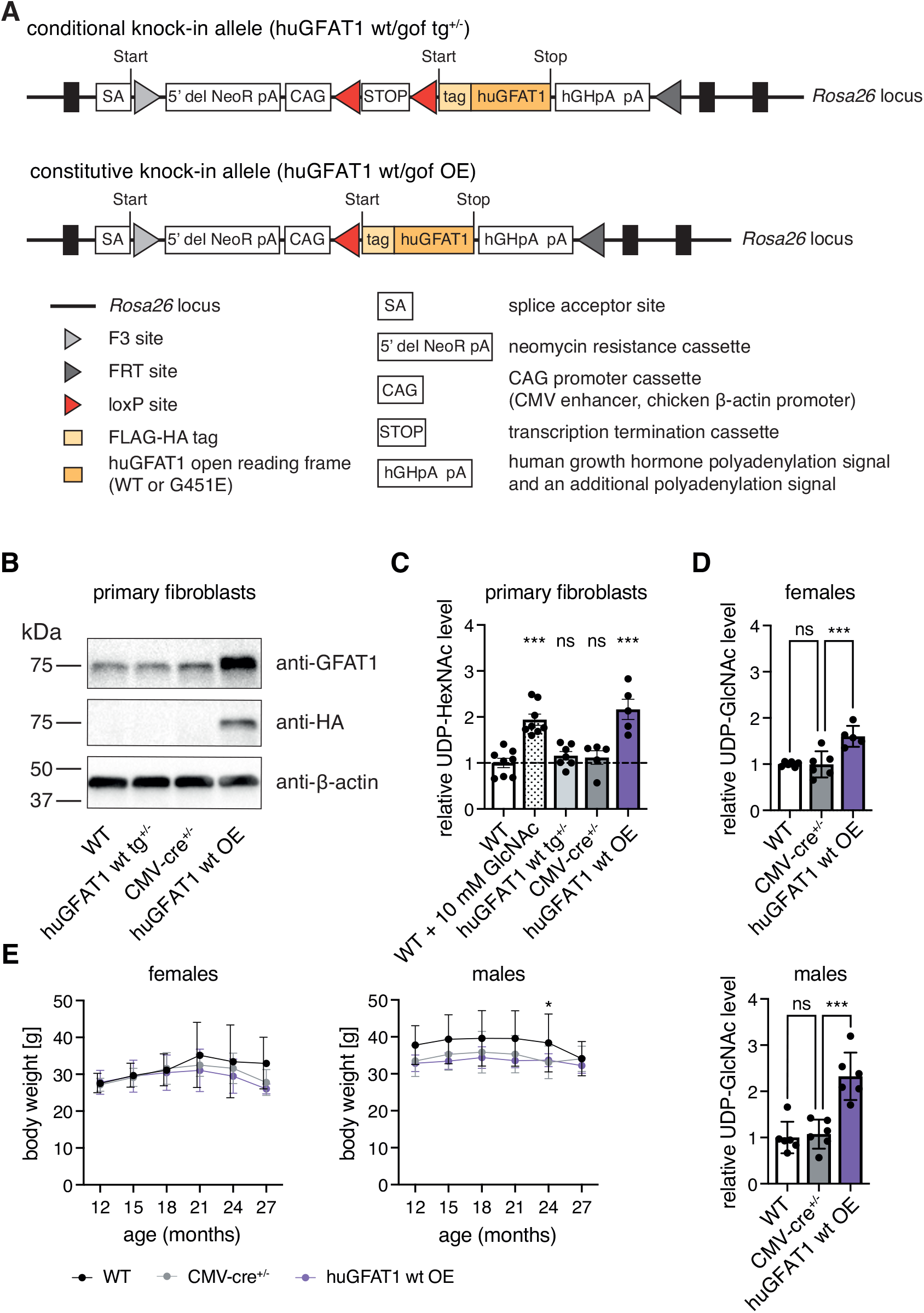
HBP activation by huGFAT1 wt OE does not influence body weight in mice. (A) Schematic representation of the transgene. The expression cassette was inserted in the Rosa26 locus (conditional knock-in allele). Upon cre-mediated deletion of the transcription termination cassette, FLAG-HA tagged human GFAT1 is expressed under the control of the chicken β-actin promoter (constitutive knock-in allele). (B) Western blot analysis of GFAT1 and HA expression in primary fibroblasts isolated from newborn mice (n=1). β-actin was used as loading control. (C) Relative UDP-HexNAc levels in primary fibroblasts. (D) Relative UDP-GlcNAc levels in hemibrain isolated from 3 months old control and huGFAT1 wt OE mice of both sexes. (C-D) Data are presented as mean ± SEM (n≥5). One-way ANOVA, Dunnett’s post- test; *** p<0.001; ns: not significant. (E) Body weight of control and huGFAT1 wt OE mice of both sexes from 12 to 27 months of age. Data are presented as mean ± SD (n≥2). Two-way ANOVA, Dunnett’s post-test. Statistical significance was calculated compared to CMV-cre^+/-^ mice at each time point; only significant changes are indicated. * p<0.05

In addition, we generated huGFAT1 gain-of-function (gof) transgenic mice using the same strategy employed for the overexpression of wt huGFAT1. The GFAT-1 G451E substitution is associated with *C. elegans* longevity (Denzel et al. 2014) and confers a drastically reduced sensitivity to UDP-GlcNAc feedback inhibition (Ruegenberg et al. 2020). huGFAT1 G451E overexpression resulted in elevated levels of both UDP-GlcNAc and UDP-GalNAc in the brains of 3 months old male and female mice (Figure S3b-c). Body weight was not affected by huGFAT1 gof OE between 12 and 27 months of age compared to CMV-cre^+/-^ mice in both sexes (Figure S3d). Together, these data suggest successful genetic HBP activation without detrimental effects on overall health as assessed by body weight analysis in huGFAT1 wt and gof OE mice. As we observed no relevant differences between huGFAT1 wt or G451E gof overexpression, we focused the remainder of the analyses on the huGFAT1 wt OE mice.

### HBP activation by huGFAT1 WT OE does not influence coordination, neuromuscular function, memory, or lifespan

To study coordination and basic neuromuscular function upon genetic HBP activation, experiments using rotarod and treadmill were performed and grip strength was measured using all genotypes at three different time points (3-4, 15-16, and 21-22 months of age). In line with the results using dietary GlcNAc supplementation, genetic HBP activation did not affect grip strength (Figure 5a, Figure S4a), rotarod performance (Figure 5b), or forced maximal endurance on a treadmill (Figure 5c). To assess the effect of genetic HBP activation on survival, lifespan experiments with transgenic mice of both sexes were performed. Survival of the huGFAT1 wt or gof OE mice was indistinguishable from the corresponding genetic controls (Figure 5d, Figure S4b).

**Figure 5:**
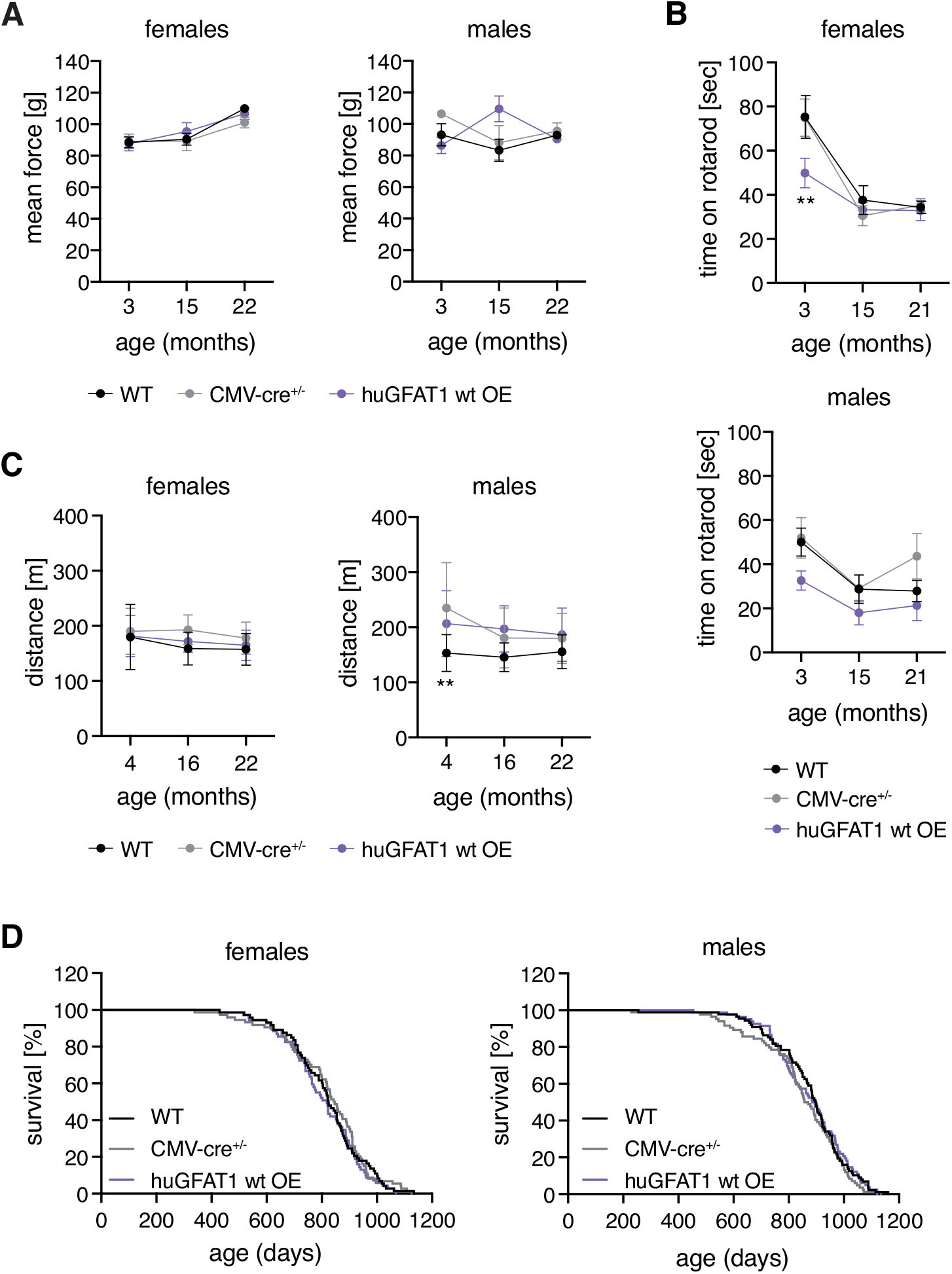
HBP activation by huGFAT1 wt OE does not affect fitness of mice. (A) Mean force measured in a grip strength test with two paws of control and huGFAT1 wt OE mice of both sexes at 3, 15 and 22 months of age. (B) Maximal time on a rotarod of control and huGFAT1 wt OE mice of both sexes at 3, 15 and 21 months of age. (A-B) Data are presented as mean ± SEM (n≥2). (C) Maximal distance on a treadmill of control and huGFAT1 wt OE mice of both sexes at 4, 16 and 22 months of age. Data are presented as mean ± SD (n≥4). (A-C) Two-way ANOVA, Dunnett’s post-test. Statistical significance was calculated compared to CMV-cre^+/-^ mice at each time point; only significant changes are indicated. ** p<0.01 (D) Lifespan analysis of control and huGFAT1 wt OE mice of both sexes (females: n≥68; males: n≥80). Survival of WT and CMV-cre^+/-^ mice is also shown in Figure S4b.

We assessed both general locomotor activity and exploratory behavior in the open field test at 3, 15 and 21 months of age in both sexes. Activity and speed of the mice was influenced by cre expression, since CMV-cre^+/-^ and huGFAT1 wt OE mice moved more and faster compared to WT controls (Figure 6a, Figure S5a). This effect was more pronounced in male mice than in females. Nevertheless, overall, genetic HBP activation had no effect on spontaneous locomotor activity. Exploratory behavior as measured by the percentage of distance spend in the center of the open field was similar in all genotypes (Figure 6b). Memory and learning were tested in both Y maze and Morris water maze. In the Y maze, alternations, i.e. how often a mouse chooses to explore a new arm over the same arm, did not change among the different genotypes, suggesting no differences in exploratory behavior and spatial working memory (Figure 6c). The distance covered in the Y maze was comparable across all genotypes in females, while it was increased in males upon cre expression (Figure S5b). In the Morris water maze, there was no difference in learning in 4 months old mice, since all genotypes of both sexes showed a similar improvement regarding distance and latency during the training sessions (Figure S5c-d). The speed of the mice was slightly decreased on day 5 compared to day 1, however, this effect was similar in all genotypes (Figure S5e). Memory, which was tested by the removal of the platform, was unchanged, since the time spent in the target quadrant (Figure 6d), the the latency of the first platform visit (Figure 6e), and the number of platform visits (Figure 6f) were similar in all genotypes. Altogether, and in contrast to dietary GlcNAc supplementation, genetic HBP activation did not improve memory in mice. Overall, results obtained using the genetic model support the conclusion that HBP activation does not have adverse effects in mice. While lifespan was not affected in any of the HBP activation regimens tested, GlcNAc supplementation improved memory in young male mice.

**Figure 6:**
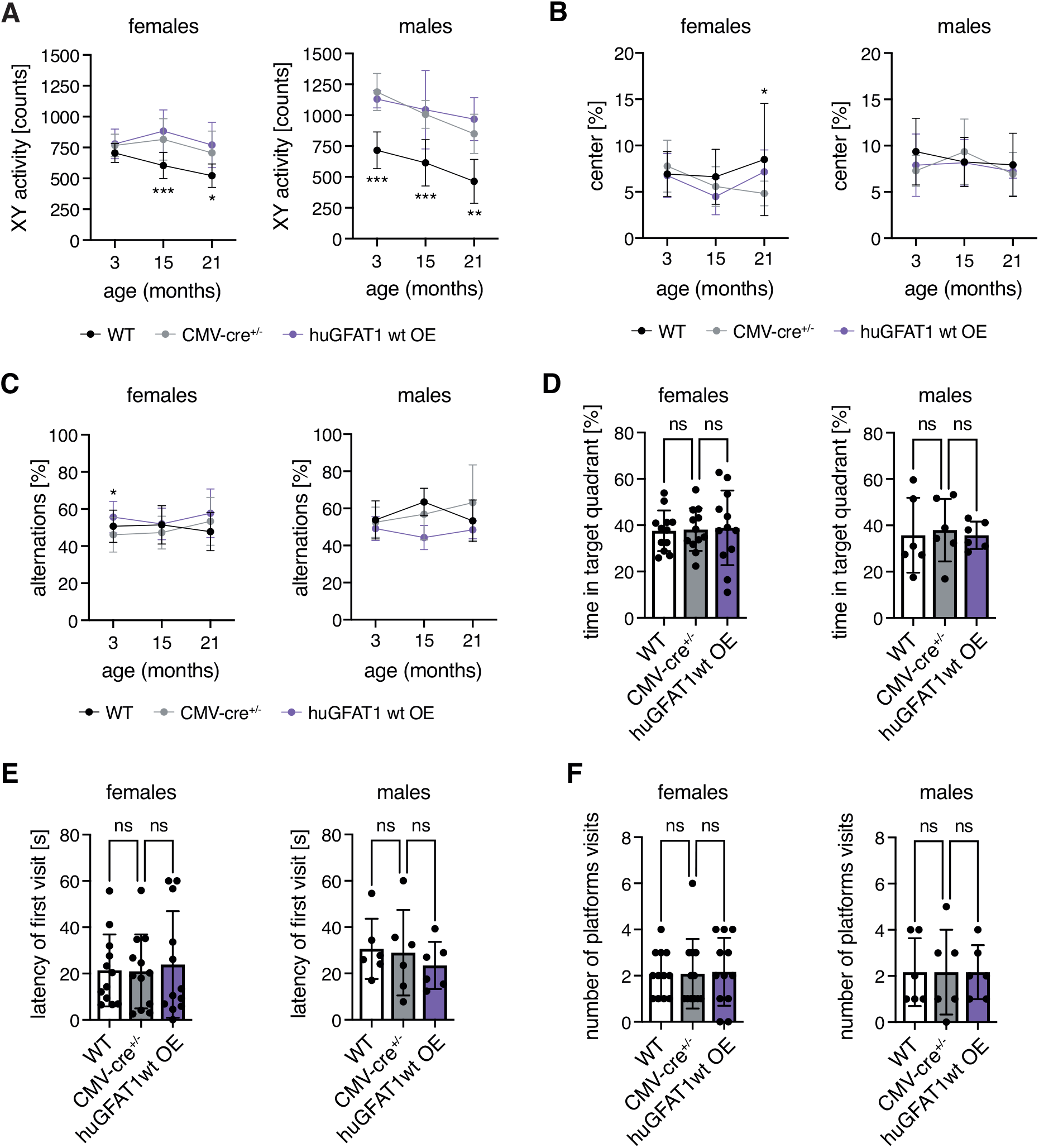
HBP activation by huGFAT1 wt OE does not influence spontaneous locomotor and exploratory behavior or memory of mice. (A) XY activity measured in the open field test of control and huGFAT1 wt OE mice of both sexes at 3, 15 and 21 months of age. (B) Percent of distance spent in the center of the open field of control and huGFAT1 wt OE mice of both sexes at 3, 15 and 21 months of age. (C) Percent of alternations measured in the Y maze test of control and huGFAT1 wt OE mice of both sexes at 3, 15 and 21 months of age. (A-C) Data are presented as mean ± SD (n≥4). Two-way ANOVA, Dunnett’s post-test. Statistical significance was calculated compared to CMV-cre^+/-^ mice at each time point; only significant changes are indicated. *** p<0.001; ** p<0.01; * p<0.05 (D) Percent of time spend in the target quadrant, (E) Latency of the first platform visit, and (F) Number of platform visits upon removal of the hidden platform in the Morris water maze test of control and huGFAT1 wt OE mice of both sexes at 4 months of age. (D-F) Data are presented as mean ± SD (n≥6). One-way ANOVA, Dunnett’s post-test; ns: not significant.

## Discussion

In this study, we delineate the effect of HBP activation on general behavior, memory, and survival in mice. Importantly, we show that dietary GlcNAc supplementation or genetic GFAT1 overexpression do not have negative side effects on the overall health of mice. While fitness, locomotion, behavior, and survival were not changed by HBP activation, we observed memory improvement in young male mice fed with GlcNAc, suggesting a beneficial effect of HBP activation in mammals.

GlcNAc is rapidly absorbed and enters the systemic circulation when fed via the drinking water: experiments using ^13^C6-GlcNAc indicate that GlcNAc peaks in the serum after 30 min of gavage and is cleared from the circulation after 2-3 h; by this time, UDP-^13^C6-GlcNAc can be detected in liver, kidney, and spleen (Ryczko et al. 2016). ^13^C6-GlcNAc, orally supplemented at a concentration of 0.1%, has been shown to cross the blood brain barrier after feeding 8 weeks old mice for 3 days: LC-MS/MS analysis identified UDP-[U^13^C]-HexNAc in the brain of adult females (Sy et al. 2020). These data indicate that GlcNAc fed via the drinking water contributes to the UDP-GlcNAc pool in brain and liver. While we could show in this (Figure 4c) and in previous studies that GlcNAc supplementation is sufficient to increase UDP-GlcNAc levels in worms and in cells (Denzel et al. 2014; Horn et al. 2020) the levels of UDP-HexNAc were not increased in the brain or liver of mice fed with 1% GlcNAc after 7 months of feeding (Figure 1d). Previously, it has been shown that GlcNAc feeding elevates hepatic UDP-HexNAc levels by around 25% (Ryczko 2016). Given this minor effect and the fast clearance of GlcNAc from the circulation, the timing of sample collection might have influenced our results: The mice were sacrificed during the day, when they drink less. Additionally, we analyzed steady-state UDP-HexNAc levels; thus, we cannot exclude elevated HBP flux upon dietary GlcNAc supplementation.

Dietary GlcNAc supplementation has previously been shown to increase body weight: post-weaning C57BL/6 male mice fed with 0.5-1.5% GlcNAc in the drinking water had a surge in body weight after around 3 months of feeding, with a concomitant increment in hepatic UDP-GlcNAc levels (Ryczko et al. 2016). In contrast, 25 months old mice fed for about 8 months with 10 g GlcN per kg of diet did not show an increase in body weight (Weimer et al. 2014). Of note, UDP-GlcNAc levels were not elevated in these animals, suggesting that body weight and UDP-GlcNAc levels correlate. In this study, we did not observe significant differences in body weight in males up to 30 months of age (Figure 1g). In contrast, females fed with 1% GlcNAc showed significantly increased body weight up to 10 months of age; later in life, the body weight was slightly, but not significantly elevated (Figure 1g). Interestingly, 9 months old females consumed more GlcNAc-containing water (Figure 1b); however, they did not display significantly elevated UDP-HexNAc levels (Figure 1d). Thus, GlcNAc intake rather than UDP-HexNAc levels correlated with body weight in this study.

HBP activation has also been linked to impaired insulin signaling: adipocytes treated with GlcN become insulin resistant (Marshall, Bacote, and Traxinger 1991) and rats infused with GlcN also exhibit insulin resistance (Virkamaki 1997), similar to GlcN- treated mice (Weimer et al. 2014). In our experiments, GlcNAc feeding did not alter the response to insulin up to 20 months of age (Figure S1c). In line with our results, although in a different animal model using pellet-based supplementation, rats exposed to 5% of GlcNAc for 1 year did not exhibit changes in the basal levels of serum glucose (Takahashi et al. 2009). Overall, these data suggest that GlcN-mediated HBP activation impairs insulin signaling, whereas dietary GlcNAc supplementation does not.

In worms, HBP activation by GlcNAc supplementation as well as GOF mutations in GFAT-1 extend lifespan (Denzel et al. 2014). In this study, the lifespan of mice fed with 1% GlcNAc was indistinguishable from controls (Figure 2e). However, based on the unaltered UDP-HexNAc levels in the brains of GlcNAc-fed animals, the HBP activity achieved here might not be sufficient to extend lifespan. Alternatively, the mechanism underlying the lifespan extension might not be conserved between worms and mammals. Importantly, GlcNAc feeding did not shorten mammalian lifespan and the overall health of the mice was not negatively affected by the treatment.

Putative LOF mutations in *GFAT1* are associated with congenital myasthenic syndrome, which is characterized by defective neuromuscular junctions (Senderek et al. 2011). Accordingly, activating the HBP pathway by GlcNAc feeding might have beneficial effects on neuromuscular function. However, we did not observe differences in coordination and fitness upon GlcNAc feeding (Figure 2a-d). While spontaneous locomotor activity and exploratory behavior were unchanged in GlcNAc-fed mice compared to controls (Figure 3a-b), GlcNAc-fed young males exhibited memory improvement in the Morris water maze (Figure 3d-e). Of note, various studies have demonstrated the contribution of myelin formation to memory consolidation and recall (Pan et al. 2020; Steadman et al. 2020). Strikingly, the HBP has a primary role in myelination: genetic or toxin-driven blockage of N-glycan branching induces demyelination, a phenotype that can be rescued by oral GlcNAc feeding (Sy et al. 2020). Furthermore, the inducible deletion of O-GlcNAc transferase -the enzyme responsible for adding GlcNAc to substrate proteins- results in learning and memory deficits (Wheatley et al. 2019). Thus, usage of the HBP’s end-product for post-translational modifications plays an important role in memory. Overall, these data suggest that the observed memory improvement in young males fed with 1% GlcNAc could be a consequence of increased myelination driven by elevated HBP activity.

Although the steady-state UDP-HexNAc levels in GlcNAc-fed mice were indistinguishable from controls (Figure 1d), we cannot exclude enhanced HBP flux. To corroborate our findings obtained with dietary GlcNAc supplementation, we additionally analyzed a model of genetic HBP activation. While ubiquitous overexpression of wt GFAT1 led to a marked increase in GFAT1 protein (Figure 4b), brain UDP-HexNAc levels were only slightly elevated in 3 months old mice (Figure 4d, Figure S3a). Moreover, the overexpression of gain-of-function GFAT1 G451E, which displays reduced sensitivity to UDP-GlcNAc feedback inhibition (Ruegenberg 2020), activated the HBP to a similar extent as wt GFAT1 OE (Figure S3b-c). Potentially, the N-terminal tag interfered with GFAT1 activity, as previously described (Olchowy et al. 2006). Nevertheless, the mice had around 2-fold increased UDP-HexNAc levels, demonstrating successful HBP activation.

GFAT1 overexpression in different tissues (namely skeletal muscle, fat, liver, and β cells) leads to disease-like metabolic states such as insulin resistance, obesity, hyperlipidemia, and impaired glucose metabolism (Hebert et al. 1996; McClain et al. 2002; Veerababu et al. 2000). While GFAT1 overexpression in liver has been associated with overweight (Veerababu et al. 2000), our results indicate that both GFAT1-overexpressing mice and controls have similar body weight profiles over the lifetime of the animals (Figure 4e and S3d). The impact of GFAT1 overexpression on coordination and neuromuscular function has never been approached before; strikingly, our results indicate that GFAT1 overexpression did not affect fitness, spontaneous activity, or exploratory behavior (Figure 5a-c, Figure S4a and Figure 6a- b). In contrast to GlcNAc feeding, GFAT1 overexpression did not enhance learning and memory formation compared to controls (Figure 6c-f and Figure S5c-e). These data suggest that the GlcNAc-mediated effect on memory might be independent of HBP activity. Of note, GlcN supplementation has been previously shown to impair glucose metabolism and to promote mitochondrial biogenesis via activation of AMPK (Weimer et al. 2014). Therefore, investigating the role of these mechanism in the GlcNAc-mediated memory improvement would be of interest in the future.

Finally, neither the overexpression of wt GFAT1 nor the G451E mutant had an impact on mouse survival (Figure 5d and Figure S4b). In contrast to GlcNAc-fed mice (Figure 1d), the transgenic mice did show increased UDP-GlcNAc levels (Figure 4d and Figure S3a-c). Thus, while HBP activation results in longevity in *C. elegans* (Denzel et al. 2014), it was not sufficient to extend lifespan in mice. In this study, either HBP activation did not reach sufficient UDP-GlcNAc levels for lifespan extension or the lifespan-modulating effects of HBP activation in the worm are caused by non- conserved mechanisms.

In sum, this study dissects the effects of dietary GlcNAc supplementation and genetic HBP activation on murine health, behavior, memory, and lifespan. Despite being linked to diabetes, obesity, and cancer, these interventions did not have adverse effects in mice. Instead, we show that GlcNAc feeding increases memory formation of young male mice. Of note, this effect might be independent of HBP activity, since we did not observe changes in memory upon genetic HBP activation. In the future, HBP- independent effects of GlcNAc supplementation should be tested in the context of learning and memory.

## Experimental procedures

### Mouse husbandry

Animals were housed on a 12:12 h light:dark cycle with *ad libitum* access to food under pathogen-free conditions in individually ventilated cages. All animals were kept in C57BL/6J background. Animal care and experimental procedures were in accordance with the institutional and governmental guidelines.

### GlcNAc feeding

Mice were fed with control water or water containing 1% GlcNAc (w/v) beginning at 8 weeks of age. GlcNAc was provided by Wellesley Therapeutics Inc., Toronto, Canada.

### Generation of transgenic mice

Generation of transgenic GFAT1 mice was performed by Taconic Biosciences (Cologne, Germany). An expression cassette was inserted in the *Rosa26* locus using recombination-mediated cassette exchange in embryonic stem cells. The cassette encodes a loxP-flanked transcription termination cassette upstream of the human GFAT1 (huGFAT1) open reading frame (conditional knock-in allele, huGFAT1 wt/gof tg^+/-^). Upon cre-mediated deletion of the transcription termination cassette, huGAFT1 is expressed under the control of the chicken β-actin promoter, resulting in its overexpression (constitutive knock-in allele, huGFAT1 wt/gof OE). huGFAT1 is N-terminally tagged with FLAG-HA (HA: hemagglutinin; for further information, see Figure 4). Mice expressing the cre recombinase under the control of the human cytomegalovirus minimal promoter (CMV-cre^+/-^) were purchased from Charles River Laboratories (Sulzfeld, Germany).

### Breeding of transgenic mice

For breeding, huGFAT1 wt/gof tg^+/-^ males were crossed with transgenic CMV-cre^+/-^ females. Both transgenes were maintained in a heterozygous state. Since the CMV promoter is active before implantation during early embryogenesis (Schwenk, Baron, and Rajewsky 1995), animals expressing the cre recombinase and carrying the GFAT1 transgene were considered to overexpress huGFAT1 in all tissues. The offspring was genotyped as described below.

### Tissue collection

Mice were sacrificed by cervical dislocation. Brains from 3 months old (genetic model) or 9 months old (GlcNAc feeding) male and female mice were dissected. Cerebral hemispheres were snap frozen in liquid nitrogen and stored at -80°C until further use.

### Lifespan analysis

Lifespan analysis was performed with 69 female and 82 male mice per genotype/condition. The general health of the mice was monitored regularly and the mice were euthanized when necessary, according to a pre-defined score sheet. The lifespan analysis comprises all mice that died a natural death or were euthanized. The body weight of the mice was analyzed every other week (GlcNAc feeding) or every 3 months (huGFAT1 wt/gof OE). To exclude an effect on lifespan, no other experiments were performed with these mice.

### Metabolic cages

Indirect metabolic analyses were performed in singly housed 3 months and 9 months old mice for 48 h using metabolic cages (Phenomaster, TSE Systems). Mice were habituated to the cages for 24 h before the measurements. Analysis of the air before and after passing the cages allowed calculation of oxygen consumption and carbon dioxide production. These values were used to determine the metabolic respiratory quotient of these mice. Additionally, the spontaneous locomotor activity, as well as food and water consumption of the mice were monitored.

### Glucose and insulin tolerance tests

Glucose and insulin tolerance was determined in 4 months, 10 months, and 20 months old mice upon GlcNAc feeding. For the glucose tolerance test, mice were fasted for 16 h with full access to drinking water. 2 g glucose per kg body weight were injected intraperitoneally. To monitor blood glucose concentration, a blood sample was taken before and 15, 30, 60, and 120 min after the glucose injection. For the insulin tolerance test, 0.75 U insulin per kg body weight were injected intraperitoneally. To monitor blood glucose concentration, a blood sample was taken before and 15, 30, and 60 min after the insulin injection. Blood glucose concentration was measured using an automatic glucose monitor (Accu-Check Aviva, Roche).

### Grip strength measurements

Grip strength measurements were performed at 6 months of age (GlcNAc feeding) or with 3, 15, and 22 months of age (huGFAT1 wt OE). The mouse is holding a trapeze while being pulled backward until the pulling force is bigger than the grip strength of the mouse (2 paws). The maximal grip strength of the mouse is recorded automatically. Alternatively, the mouse is placed on a grid and the grasping applied by the mouse while being pulled backwards is measured (4 paws). The tests were repeated 5 times per mouse per day and the mean of the 5 trials is plotted.

### Open field

Open field analyses were performed with 6 months of age (GlcNAc feeding) or with 3, 15, and 21 months of age (huGFAT1 wt OE). The mice were placed in a 50x50x40 cm big box for 10 min. The spontaneous locomotor and explorative activity, as well as the speed were analyzed by tracking the movement of the mice.

### Rotarod

Rotarod analyses were performed at 6 months of age (GlcNAc feeding) or with 3, 15, and 21 months of age (huGFAT1 wt OE). The rotation speed of the rod constantly increased from 5 to 40 U/min within 5 min. The time until the mice fell of the rod was measured. The experiment was performed twice per day on two or four consecutive days for GlcNAc feeding and huGFAT1 wt OE mice, respectively. The average time on the rod of the two runs on the last day of the experiment is plotted.

### Treadmill

Treadmill analyses were performed at 6 months of age (GlcNAc feeding) or with 4, 16, and 22 months of age (huGFAT1 wt OE). After an adaption phase of 5 min, the treadmill was started with a speed of 0.1 m/s. This speed was maintained for 10 min before it constantly increased to 1.3 m/s within 60 min. An electric shock of 0.3 mA was given whenever the mouse stayed at the end of the treadmill for more than 2 sec. The experiment was stopped when a mouse received three consecutive shocks. The cumulative distance was analyzed for each mouse.

### Y maze

Y maze analyses were performed with 3, 15, and 21 months of age (huGFAT1 wt OE). The mouse is placed in one arm of the Y maze and can explore the environment for 5 min. The activity of the mouse is tracked. Distance and alternations are measured for each mouse. The alterations describe how often a mouse chooses to explore a new region over the same region.

### Water maze

Water maze analyses were performed with 4 months of age (GlcNAc feeding and huGFAT1 wt OE). The mice were placed in a basin filled with stained water, in which a platform was hidden 1-2 cm below the surface of the water. Since the mice want to avoid swimming, they will learn to find the hidden platform during training sessions. Here, the mice were trained for five consecutive days on which they performed four trials per day, with a maximum duration of 60 s per trial. The average of the four trials is plotted for each day. After the training on day 5, the platform was removed. During this final test, the memory of the mice was analyzed. The mice were placed in the basin for 60 s and their preference for the quadrant which previously contained the platform, as well as the latency of the first visit and the number of visits of the former platform was analyzed.

### Isolation of mouse genomic DNA from ear clips

Ear clips were taken by the Comparative Biology Facility at the Max Planck Institute for Biology of Ageing (Cologne, Germany) at weaning age (3-4 weeks of age) and stored at -20°C until use. 150 µl ddH2O and 150 µl directPCR Tail Lysis reagent (Peqlab) were mixed with 3 µl proteinase K (20 mg/ml in 25 mM Tris-HCl, 5 mM Ca2Cl, pH 8.0, Sigma-Aldrich). This mixture was applied to the ear clips, which were then incubated at 56°C overnight (maximum 16 h) shaking at 300 rpm. Proteinase K was inactivated at 85°C for 45 min without shaking. The lysis reaction (2 µl) was used for PCR without further processing. For genotyping of newborn pups, the tail tip was lysed in 300 µl directPCR Tail Lysis reagent containing 3 µl proteinase K.

### Genotyping PCR

For genotyping of mouse genomic DNA DreamTaq DNA polymerase (ThermoFisher Scientific) was used. The presence of the cre transgene was determined using Cre_fwd (GCCAGCTAAACATGCTTCATC) and Cre_rev (ATTGCCCCTGTTTCACTATCC). The primer targeting huGFAT1 (huGFAT1_fwd CGGTGGAGGTTACCCATACG; huGFAT1_rev CGAGCTTGGCAATTGTCTCTG) detected presence of the transgene without distinction of zygosity. The product amplified using oIMR7338 and oIMR7339 served as internal control for huGFAT1 and cre transgene detection. Using the huGFAT1_R26 PCR (3224_35 TTGGGTCCACTCAGTAGATGC; 1114_1 CTCTTCCCTCGTGATCTGCAACTCC; 1114_2 CATGTCTTTAATCTACCTCGATGG ), the wildtype Rosa26 locus and the targeted Rosa26 locus could be distinguished.

### Isolation and maintenance of primary fibroblasts

For fibroblast isolation newborn mice (P0-P3, both sexes) were sacrificed by decapitation. The corpus was incubated in 50% betaisodona/PBS (Mundipharma GmbH) for 30 min at 4°C before being washed in different solutions for 2 min each: PBS (ThermoFisher Scientific), 0.1% octenidin in ddH2O (Serva Electrophoresis), PBS, 70% ethanol, PBS, antibiotic-antifungal-solution in PBS (ThermoFisher Scientific). Tail and legs were removed and the tail tip was used for genotyping. Complete skin was separated from the body and incubated in 2 ml dispase II solution (5 mg/ml in 50 mM HEPES/KOH pH 7.4, 150 mM NaCl; Sigma-Aldrich) over night at 4°C. The epidermis was separated from the dermis as a sheet. The dermis was minced into small pieces using scalpels and transferred to a falcon tube containing collagenase (400 U/ml in 50 mM Tris base, 5 mM CaCl2, pH 7.4; Sigma-Aldrich). The samples were incubated at 37°C for 1.5 h and mixed regularly. Next, the suspension was filtered through a 70 µm cell strainer, which was washed with DMEM afterwards. The cells were centrifuged for 10 min at 1000 rpm. The pellet was resuspended in DMEM (4.5 g/L glucose, 10% fetal bovine serum and penicillin/streptavidin; all ThermoFisher Scientific) and the cells were seeded on non-coated tissue culture plates. The cells were grown at 37°C in 5% CO2.

### Western Blot analysis

Protein concentration of cell lysates was determined using the Pierce^TM^ BCA protein assay kit according to manufacturer’s instructions (ThermoFisher Scientific). Samples were subsequently subjected to SDS-PAGE and blotted on a nitrocellulose membrane. The following antibodies were used in 5% low-fat milk (Carl Roth) or 1% bovine serum albumin (BSA; Carl Roth) in TBS-Tween buffer (25 mM Tris base, 150 mM NaCl, 2 mM KCl, pH 7.4; 0.05% Tween-20 (w/v)) over night at 4°C: GFPT1 (rabbit, Abcam, EPR4854, 1:1.000 in BSA), hemagglutinin (HA; rat, Roche Diagnostics GmbH, 3F10, 1:1.000 in milk), β-actin (mouse, Cell Signaling Technologies, 8H10D10, 1:25.000 in milk). After incubation with HRP-conjugated secondary antibody (Invitrogen, 1:5000), the blot was developed using ECL solution (Merck Millipore). Bands were detected on a ChemiDoc MP Imaging System (Bio-Rad Laboratories).

### Metabolite analysis

#### Determination of UDP-HexNAc levels

Brains were collected from 9 months old mice, cut in half, and snap frozen in liquid nitrogen. For metabolite extraction, 250 µl ddH2O were added to the hemibrains and the tissue was disrupted using a dounce homogenizer. The samples were subjected to four freeze/ thaw cycles (liquid nitrogen/ 37°C water bath). Next, the protein concentration was measured using the Pierce^TM^ BCA protein assay kit (ThermoFisher Scientific). 200 µl with a protein concentration of 1 µg/µl were mixed with 1 ml chloroform:methanol (1:2) and incubated on a nutator mixer for 1 h at RT. After centrifugation for 5 min at full speed, the supernatant was transferred to a glass vial. The liquid was evaporated in an EZ-2 Plus Genevac centrifuge evaporator (SP Scientific) with the following settings: time to final stage 15 min, final stage time 4 h, low boiling point mixture. After evaporation, the samples were stored at -20°C until further use.

Absolute UDP-HexNAc levels were determined using an Acquity UPLC connected to a Xevo TQ Mass Spectrometer (both Waters) and normalized to total protein content. The measurements and subsequent analysis were performed as previously described (Denzel et al. 2014).

#### Determination of UDP-GlcNAc and UDP-GalNAc levels

Brains were collected from 3 months old mice, cut in half, and snap frozen in liquid nitrogen. Tissue was disrupted using a TissueLyser II (Qiagen) at 20-25 Hz. The powder was transferred to a fresh tube and subjected to metabolite extraction.

Metabolite extraction was performed using 80% methanol. After vortexing, the samples were incubated at -20°C for 30 min. Afterwards, samples were incubated on an orbital mixer at 5°C for 30 min. The samples were centrifuged for 5 min at full speed and 4°C. The supernatant was transferred to a fresh tube and the pellet was used for protein extraction with 0.5% SDS. The supernatant was evaporated in a SpeedVac concentrator at 25°C.

The metabolite analysis was conducted using a Dionex ICS-5000 anion exchange chromatography (ThermoFisher Scientific). Separation was performed with a Dionex Ionpac AS11-HC column (2 mm x 250 mm, 4 μm particle size, Thermo Fisher) at 30°C. A guard column, Dionex Ionpac AG11-HC b (2 mm x 50 mm, 4 μm particle size, ThermoFisher Scientific), was placed before the separation column. The eluent (KOH) was generated by a KOH cartridge using ddH2O. A gradient was used for the separation at a flow rate of 0.380 ml/min: 0-8 min 30 mM KOH, 8-12 min 35-100 mM KOH, 12-15 min 100 mM KOH, 15-19 min 30 mM KOH. A Dionex suppressor AERS 500 (2 mm) was used for the exchange of KOH and operated with 95 mA at 17°C. The suppressor pump flow was set to 0.6 ml/min. Samples were diluted in ddH2O and injected from a tempered autosampler (8°C) using full loop mode (10 µl). The Dionex ICS-5000 was connected to a XevoTM TQ mass spectrometer (Waters) and operated in negative ESI MRM (multi reaction monitoring) mode. The source temperature was set to 150°C, the desolvation temperature was set to 350°C and desolvation gas was set to 650 l/h, while cone gas was set to 50 l/h. The MRM transition 606.10 → 158.80 was used for quantification of UDP-GlcNAc and UDP-GalNAc. An external standard calibration curve was prepared from 50 to 1000 ng/ml UDP-GlcNAc and UDP-GalNAc. Data were analyzed using the MassLynx and TargetLynx software (Waters).

## Acknowledgments

We thank all M.S.D. laboratory members for lively and helpful discussions. We thank the Metabolomics Core Facility and the Comparative Biology Facility at the Max Planck Institute for Biology of Ageing. We thank Maribel Schönewolff, Emanuel Bruckisch, and Klara Schilling for their support with mouse genotyping. K.A. was supported by the Cologne Graduate School for Ageing Research. M.S.D. was supported by ERC-StG 640254 and ERC-PoC 768524, by the Deutsche Forschungsgemeinschaft (DFG, German Research Foundation) – Projektnummer 73111208 - SFB 829, and by the Max Planck Society.

## Authors’ contributions

K.A, M.D.H., A.M., and M.S.D. designed the research. K.A. and M.D.H. wrote the manuscript. M.P. and A.M. performed mouse phenotyping. K.A. performed all other experiments.

## Supplementary figures

**Figure S1:**
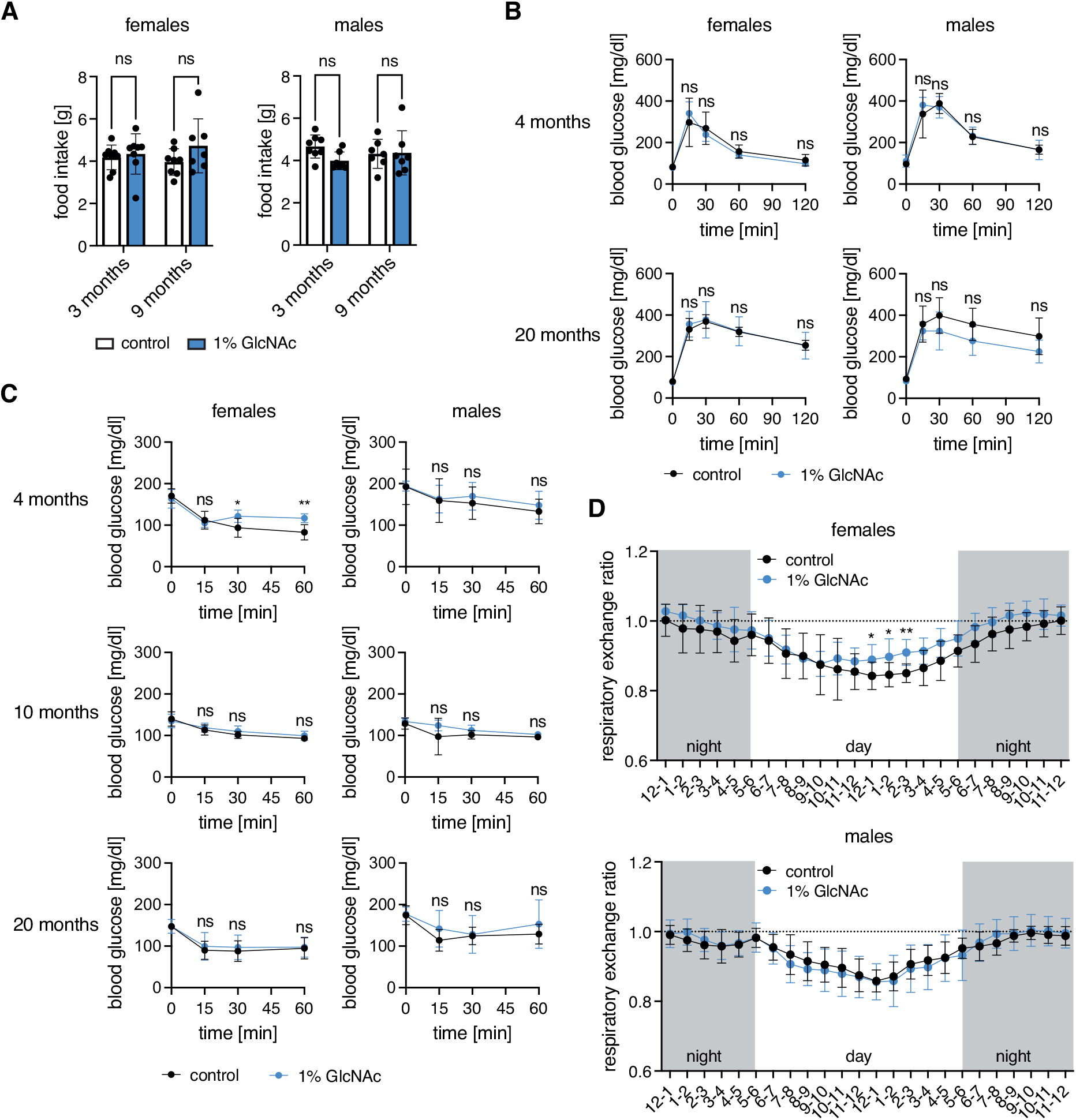
GlcNAc supplementation does not influence food intake or insulin tolerance in mice. (A) Food intake of control (white) and GlcNAc-treated mice (blue) of both sexes at 3 and 9 months of age measured in the metabolic cages. Data are presented as mean ± SD (n≥7). Two-way ANOVA, Tukey’s post-test; ns: not significant (B) Blood glucose concentration at 0 (fasting), 15, 30, 60, and 120 min after intraperitoneal injection of glucose solution (2 g/kg body weight) of control (black) and GlcNAc-treated mice (blue) of both sexes at 4 and 20 months of age. (C) Blood glucose concentration before (0), and 15, 30, and 60 min after intraperitoneal injection of insulin (75 U/kg body weight) of control (black) and GlcNAc-treated mice (blue) of both sexes at 4, 10, and 20 months of age. (B-C) Data are presented as mean ± SD (n≥6). Multiple unpaired t-tests; ** p<0.01; * p<0.05; ns: not significant (D) Respiratory exchange ratio (CO2 production/O2 consumption) of control (black) and GlcNAc- treated mice (blue) of both sexes during the day and at night (gray) at 9 months of age measured in the metabolic cages. Data are presented as mean ± SD (n≥7). Multiple unpaired t-tests; ** p<0.01; * p<0.05; only significant changes are indicated.

**Figure S2:**
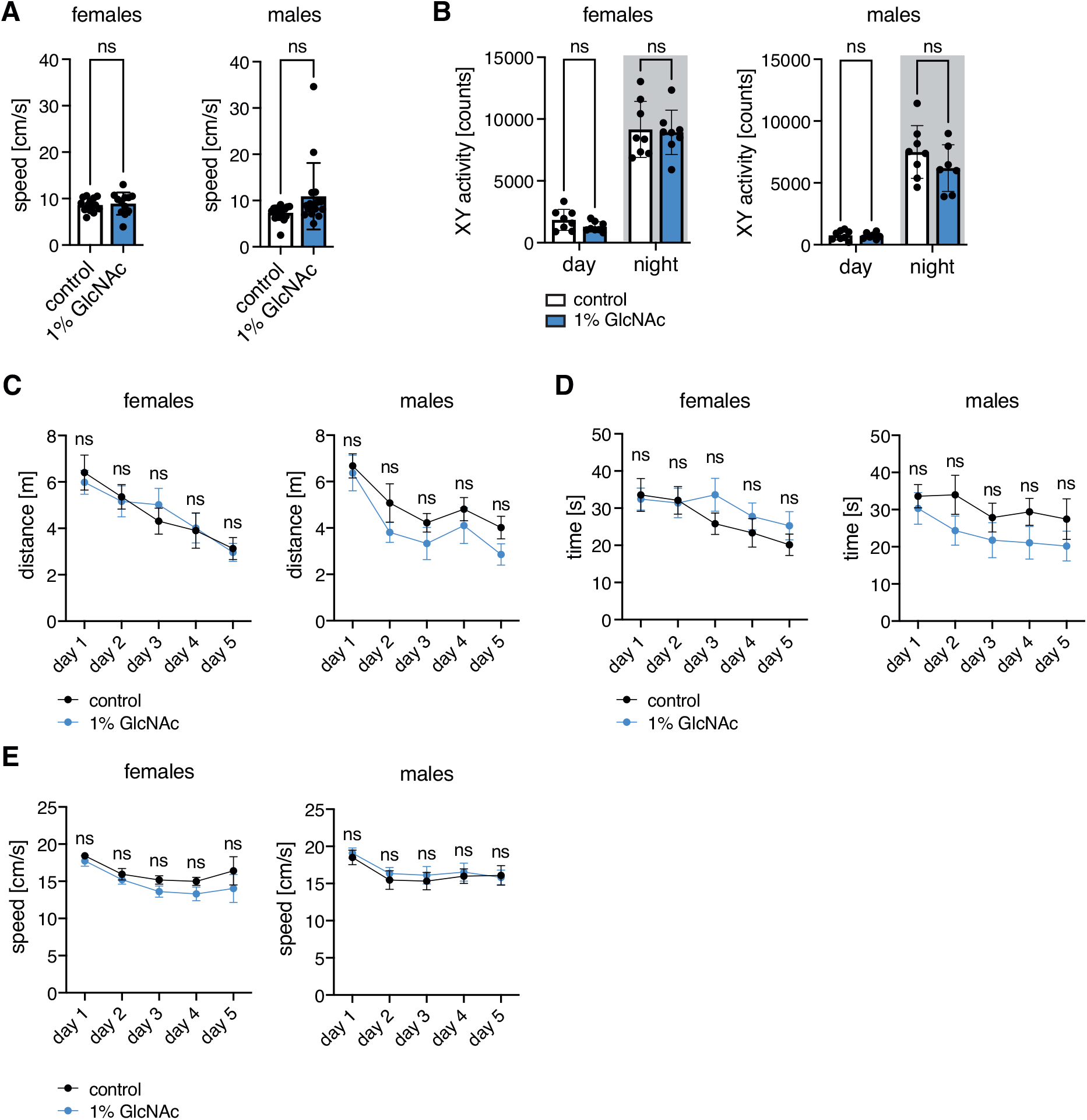
GlcNAc supplementation does not influence spontaneous locomotor behavior or learning in mice. (A) Speed of control (white) and GlcNAc-treated mice (blue) of both sexes at 6 months of age measured in the open field test. Data are presented as mean ± SD (n≥12). Unpaired t-test; ns: not significant (B) XY activity of control (white) and GlcNAc-treated mice (blue) of both sexes during the day and at night (gray) at 3 months of age measured in the metabolic cages. Data are presented as mean ± SD (n≥7). Two-way ANOVA, Tukey’s post-test; ns: not significant. (C) Distance and (D) Time until the mice reached the hidden platform, and (E) Swimming speed of control (black) and GlcNAc-treated mice (blue) of both sexes at 4 months of age during the Morris water maze training. (C-E) Data are presented as mean ± SEM. (n=12). The mean of four trials per day is plotted. Multiple unpaired t-tests; ns: not significant.

**Figure S3:**
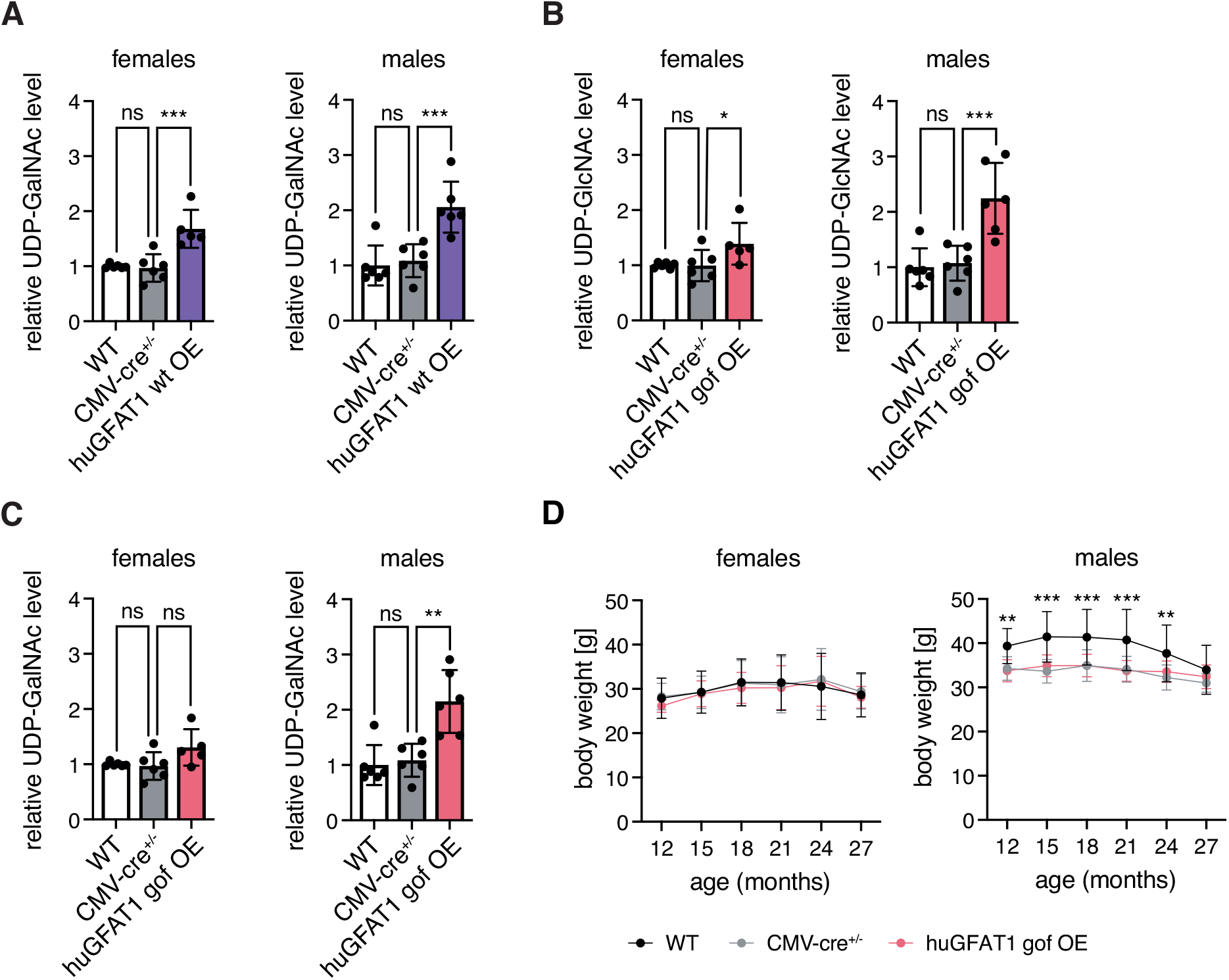
HBP activation by huGFAT1 wt/gof OE in mice. (A) Relative UDP-GalNAc levels in hemibrain isolated from 3 months old control and huGFAT1 wt OE mice of both sexes. (B) Relative UDP-GlcNAc levels in hemibrain isolated from 3 months old control and huGFAT1 gof OE mice of both sexes. (C) Relative UDP-GalNAc levels in hemibrain isolated from 3 months old control and huGFAT1 gof OE mice of both sexes. (A-C) Data are presented as mean ± SEM. One-way ANOVA, Dunnett’s post-test; *** p<0.001; ** p<0.01; * p<0.05; ns: not significant. (D) Body weight of control and huGFAT1 gof OE mice of both sexes from 12 to 27 months of age. Data are presented as mean ± SD (n≥5). Two-way ANOVA, Dunnett’s post-test. Statistical significance was calculated compared to CMV-cre^+/-^ mice at each time point; only significant changes are indicated. *** p<0.001; ** p<0.01

**Figure S4:**
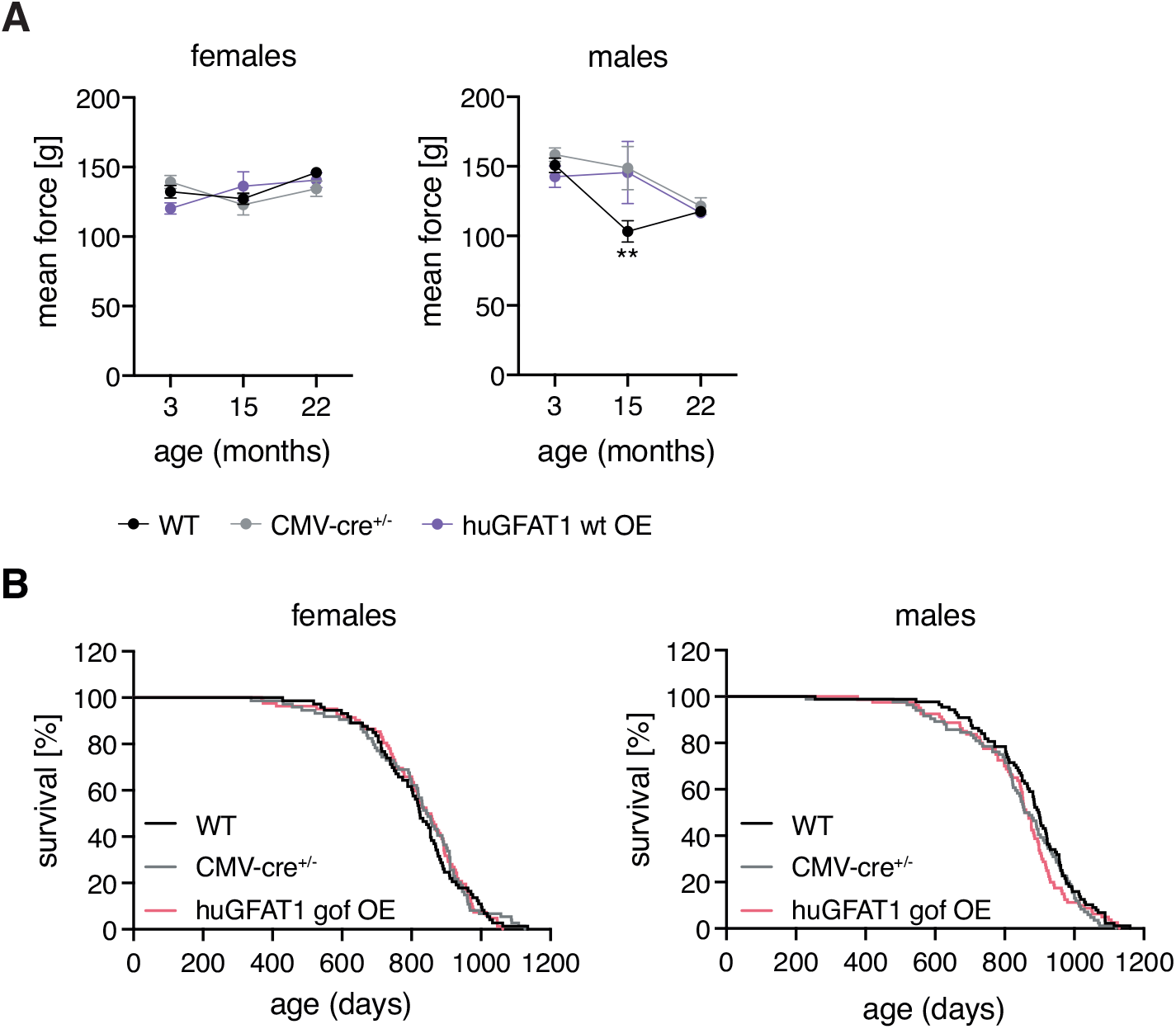
Genetic HBP activation does not affect fitness of mice. (A) Mean force measured in a grip strength test with four paws of control and huGFAT1 wt OE mice of both sexes at 3, 15 and 22 months of age. Data are presented as mean ± SD (n≥4). Two-way ANOVA, Dunnett’s post-test. Statistical significance was calculated compared to CMV-cre^+/-^ mice at each time point; only significant changes are indicated. ** p<0.01 (B) Lifespan analysis of control and huGFAT1 gof OE mice of both sexes (females: n≥68; males: n≥80). Survival of WT and CMV-cre^+/-^ mice is also shown in Figure 5d.

**Figure S5:**
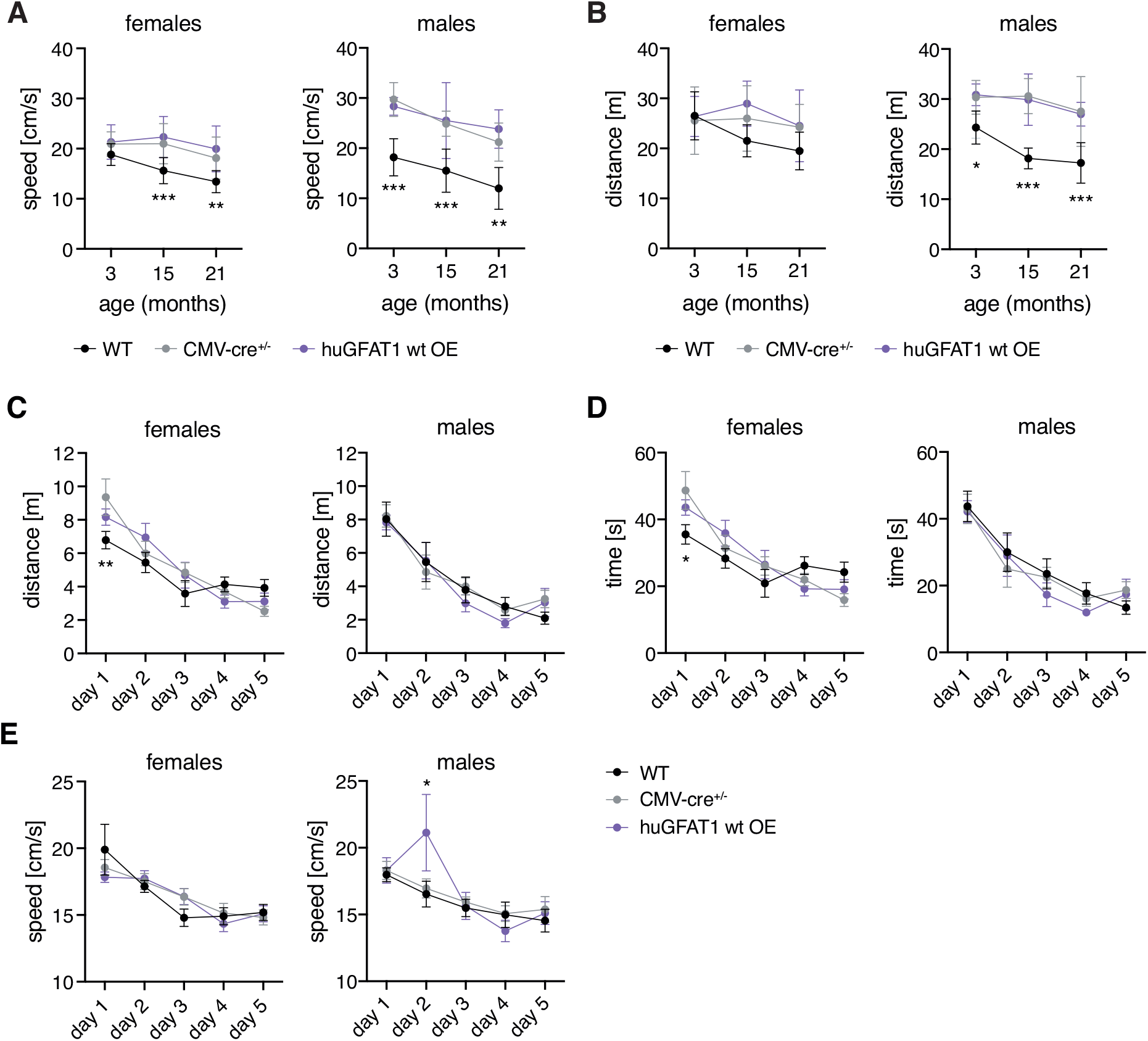
HBP activation by huGFAT1 wt OE does affect behavior or learning of mice. (A) Speed measured in the open field test of control and huGFAT1 wt OE mice of both sexes at 3, 15 and 21 months of age. (B) Distance measured in the Y maze test of control and huGFAT1 wt OE mice of both sexes at 3, 15 and 21 months of age. (A-B) Data are presented as mean ± SD (n≥4). (C) Distance and (D) Time until the mice reached the hidden platform, and (E) Swimming speed of control and huGFAT1 wt OE mice of both sexes at 4 months of age during the Morris water maze training. (C-E) Data are presented as mean ± SEM (n≥6). The mean of four trials per day is plotted. (A-E) Two-way ANOVA, Dunnett’s post-test. Statistical significance was calculated compared to CMV-cre^+/-^ mice at each time point; only significant changes are indicated. *** p<0.001; ** p<0.01; * p<0.05

